# The Molecular Mechanisms of Defensive-Grade Organic Acid Biosynthesis In Ground Beetles

**DOI:** 10.1101/2024.07.05.601757

**Authors:** Adam M. Rork, Sihang Xu, Athula Attygalle, Tanya Renner

## Abstract

Insects are known to synthesize and secrete hundreds of unique defensive chemicals, including caustic acids, pungent phenolics, and citrusy terpenes. Despite efforts to characterize the defensive chemistry of ground beetles (Coleoptera: Carabidae), our knowledge of semiochemical evolution within the family and how these compounds are biosynthesized remains limited. Few studies have demonstrated the likely biosynthetic precursors of select compounds in certain taxa, and only one has demonstrated which genes may be involved in the biosynthesis of formic acid. Here, we characterize the defensive chemistry and generate defensive gland transcriptomes for ground beetle species representing two defensive chemical classes: the formic acid producer *Platynus angustatus* and the methacrylic acid producer *Pterostichus moestus*. Through comparative transcriptome analyses, we demonstrate that co-option of distinct primary metabolic pathways may be involved in formic acid and methacrylic acid biosynthesis in the defensive glands of these taxa. These results expand our knowledge of ground beetle defensive chemistry and provide additional evidence that co-option of existing primary metabolic pathways plays a major role in the evolution of ground beetle chemical defense.

## BACKGROUND

Arthropods are well-known for not only their taxonomic diversity, but their biochemical diversity (Roth & Eisner 1962). For many species, biochemicals are the primary means of communicating ecologically relevant information with conspecifics and heterospecifics alike. Eusocial insects, such as ants, emit trail pheromones to inform nestmates about the location of resources (Attygalle & Morgan 1985). Male Lepidoptera emit sex pheromones from coremata to advertise their location to potential mates as well as repel competitors (Birch et al. 1990). Bark beetles (Scolytinae) are well known for emitting aggregation pheromones that attract swarms of conspecifics to trees for oviposition, resulting in vast quantities of larvae feeding upon the tree and overwhelming induced defenses (Symonds & Elgar 2003). Millipedes secrete toxic quinones, cyanide, and benzaldehyde in response to perturbation by organisms ignoring their aposematism (Rodriguez et al. 2018). These semiochemicals have been of interest to the scientific community for decades given their potential to explain patterns of insect evolution and behavior. Among the most well-studied are insect defensive compounds, which include small organic molecules, venoms, and complex polymers (Attygalle et al. 1993, Eisner et al. 1961, Roth & Eisner 1962, Laxme et al. 2019). The former comprise arguably the largest area of inquiry and include a wide variety of metabolites such as carboxylic acids, quinones, phenolics, terpenes, sulfides, etc. (Roth & Einser 1962, Moore & Wallbank 1968). Indeed, hundreds if not thousands of small organic defensive compounds have been described across Arthropoda, mostly from exocrine secretions. Despite our breadth of knowledge regarding the diversity of insect defensive compounds, our knowledge of their biosynthesis and evolution is wanting. Efforts have been made in recent years to fill this gap in knowledge, perhaps most notably in one of the phylum’s most diverse lineages, Adephaga (Coleoptera).

A synapomorphy of Adephaga is the unique pair of defensive glands situated in the abdomen, aptly known as pygidial glands (Fig. 1a) (Deuve 1993, Forsyth 1968, Forsyth 1970, Forsyth 1972). These glands are often composed of four gross morphological units, although some taxa have evolved additional features. These include the secretory lobes, the collecting ducts, the reservoirs, and the efferent ducts (Fig. 1b). Defensive chemical precursors are transported into secretory lobe tissue, the primary site of defensive chemical biosynthesis. Defensive compounds are then secreted from the secretory lobes into the chemically resistant, resilin-rich collecting ducts, which are transported to the reservoirs for storage (Forsyth 1972, Rork et al. 2019, Muzzi et al. 2019). When a beetle is perturbed, the reservoirs contract, propelling defensive compounds through the efferent ducts which open to the tip of the abdomen. In certain taxa, additional accessory glands have been found articulated to the efferent duct (Forsyth 1972). The efferent ducts of bombardier beetles (Brachininae, Paussinae) have evolved into sclerotized reaction chambers within which their characteristically exothermic conversion of p-hydroquinones and hydrogen peroxide to p-benzoquinones and water occurs (Aneshansley et al. 1969). In other taxa, the reservoir is divided into two lobes rather than the one typical of most taxa (Will et al. 2000).

**Figure 1:**
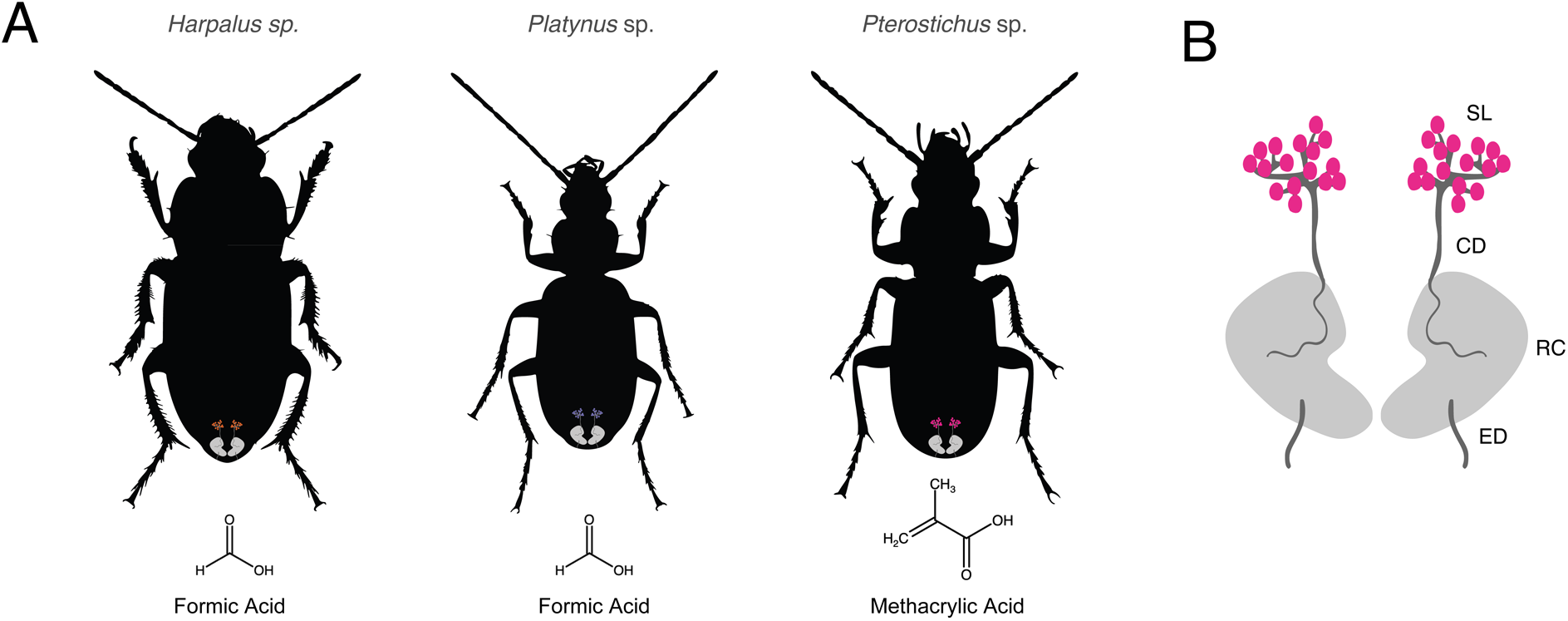
Silhouette representations of *Harpalus* sp., *Platynus* sp., and *Pterostichus* sp. with graphical representations of the pygidial glands shown at approximate locations in the abdomen (a), along with the primary defensive compound they secrete (formic or methacrylic acid). An enlarged representation of the pygidial glands is displayed in (b), noting the generalized organization consisting of the secretory lobes (SL), collecting duct (CD), reservoir chamber (RC), and efferent duct (ED). Neither glands nor species are to scale. Silhouettes are based on images of *Harpalus smaragdinus*, *Platynus livens*, and *Pterostichus macer*, all by Udo Schmidt.

Adephagan defensive chemistry is exceptionally diverse, with several hundred chemicals described and likely more left undiscovered (Rork & Renner 2018). These compounds can be categorized into broad chemical classes, including carboxylic acids, quinones, terpenes, sulfides, aromatic acids, hydrocarbons, etc. Of these, the most ubiquitous appear to be the carboxylic acids, namely methacrylic acid and formic acid (Schildknecht 1970, Kanehisa & Murase 1977, Kanehisa & Kawazu 1982, Kanehisa & Kawazu 1985, Will et al. 2000, Rork et al. 2021). Methacrylic acid is present in several adephagan subfamilies, including the Carabinae, Harpalinae, Scaratinae, and Trechinae (Moore & Wallbank 1968, Kanehisa & Murase 1977, Lečić et al. 2014). Formic acid has only been detected in the Harpalinae and Trechinae but is especially widespread in the former (Moore & Wallbank 1968, Kanehisa & Murase 1977, Moore 1979, Rossini et al. 1997, Rork et al. 2021). Also widespread are quinones, which while not being as abundant as formic or methacrylic acid, can be found in at least three subfamilies: the Brachininae, the Harpalinae, and the Paussinae (Moore & Wallbank 1968, Schildknecht et al. 1968, Kanehisa & Murase 1977). The Harpalinae biosynthesize benzoquinones directly in the secretory lobes whereas the Brachininae and Paussinae oxidize hydroquinones to benzoquinones in their reaction chambers. While often not discussed as primary defensive chemicals, aliphatic hydrocarbons, ketones, aldehydes, esters, and alcohols can be found in the pygidial gland secretions of most subfamilies (Moore 1979, Rossini et al. 1997, Will et al. 2000).

Given the most current phylogenetic hypotheses for the family Carabidae, this pattern of defensive chemical presence across the family suggests that many compounds have evolved several times independently (Vasilikopoulos et al 2021). Indeed, quinones are likely to have evolved at least three times, formic acid at least twice, terpenes twice, etc. (Moore & Brown 1979, Attygalle et al. 2009). Even the highly specialized bombardier beetle phenotype, which entails the biosynthesis of concentrated (>20%) hydrogen peroxide, has evolved twice in the family (Di Giulio et al. 2015, Muzzi et al. 2019, Vasilikopoulos et al 2021). This raises an interesting question about the nature of defensive chemical evolution in the Carabidae: where independent lineages have evolved to secrete identical defensive chemicals, do they biosynthesize these compounds using identical or distinct biochemical pathways? For example, we have previously provided transcriptomic evidence that the formic acid producer *Harpalus pensylvanicus* may biosynthesize its primary defensive compound via the folate cycle of C1 metabolism (Rork et al. 2021). However, there are several other mechanisms by which formic acid could be biosynthesized in Carabidae, including the kynurenine pathway, the methionine salvage cycle, alpha-oxidation of fatty acids, etc. (Meiser et al. 2016, Brosnan & Brosnan 2016, Badawy 2017). Perhaps the folate cycle was not only co-opted by *H. pensylvanicus*, but all formic acid-producing carabids. Alternatively, the folate cycle may have been co-opted in some taxa, the kynurenine pathway in others, etc. It may also be that some lineages biosynthesize formic acid through novel pathways. Unfortunately, aside from *H. pensylvanicus* and a few other taxa, we know very little about how carabid defensive chemicals are biosynthesized (Attygalle et al. 1991, Adachi et al. 1985, Attygalle et al. 2006, Attygalle et al. 2020, Rork et al. 2021).

Here, we aim to identify candidate genes and pathways underlying formic acid and methacrylic acid biosynthesis for two taxa belonging to the subfamily Harpalinae. We hypothesize that the formic acid producer *Platynus angustatus* (Platynini) upregulates genes involved in the folate cycle of C1 metabolism within their secretory lobes, suggesting a role in the biosynthesis of defensive-grade formic acid as in the case of *Harpalus pensylvanicus* (Rork et al. 2021). Furthermore, we hypothesize that *Pterostichus moestus* (Pterostichini) biosynthesizes methacrylic acid from L-valine and thus would upregulate genes involved in the valine catabolic pathway. This is based on previous work which has shown that the species *Carabus yaconinus* (Carabinae) and *Scarites subterraneus* (Scaratinae), biosynthesize methacrylic acid from L-valine (Adachi et al. 1985, Attygalle et al. 1991). The valine catabolic pathway does not typically lead to the biosynthesis of methacrylic acid but does generate the immediate fatty acyl-CoA precursor, methacrylyl-CoA, a probable intermediate in the acid’s biosynthesis. We also aimed to identify genes involved in defensive carboxylic acid transport in these species’ secretory lobes. Without efficient export of these compounds from gland cells into collecting duct lumens, not only would these species have no chemical stores to defend themselves with, but the glands themselves would be subject to immense physiological stress due to the buildup of cytotoxic chemicals.

## MATERIALS AND METHODS

### Beetle Collection

*Platynus angustatus* and *Pterostichus moestus* specimens were collected at Stone Valley Recreation Area in Huntingdon County, Pennsylvania throughout the summer and fall of 2019. Beetles were kept in plastic containers with coconut fiber substrate, were fed a diet of pecans and dog food (Castor & Pollux Organic Cookies, Chicken Recipe), and were provided with wet paper towels for water. Both species, especially *P. moestus*, were observed to be extraordinarily cannibalistic even when fed, and were thus isolated into individual containers.

### GC-MS Analysis of Pygidial Gland Contents

Gland exudates were collected for *Pterostichus moestus* and *Platynus angustatus* as described in previous work (Rork et al. 2021). Briefly, for each species, the hind legs of two beetles were pinched, causing them to spray into 2 ml autosampler vials containing anhydrous dichloromethane (Sigma-Aldrich). While we were able to confidently determine the sexes of *Pterostichus moestus* prior to collecting their gland exudates, and thus collect sex-specific sprays, we were unable to do so for *Platynus angustaus*. A typical external character used to determine sex in Carabidae is the size of the foretarsi, males often having broader foretarsi than females, but this dimorphism was vague-to-absent in *P. angustatus*. making reliable identification difficult without a reference or dissection. Gland extracts were analyzed on a 30 m x 0.25 mm x 0.25 μm ZB-WAX column installed in a Shimadzu 17A gas chromatograph coupled to a QP5050 mass spectrometer. The oven temperature was held at 40 °C for 4 minutes and increased at 10 °C/min to a final temperature of 240 °C, and held for 5 minutes.

### Pygidial Gland Dissections

*Platynus angustatus* and *Pterostichus moestus* specimens were dissected as described previously (Rork et al. 2021). Briefly, hind legs of beetles were pinched to induce a defensive spray response. After waiting thirty minutes, beetles were separated at the prothoracic-mesothoracic junction and were stored in RNAlater Stabilization Solution (Invitrogen) at -80 °C until dissection. Specimens were dissected under an Olympus SZX16 stereomicroscope using fine-tipped forceps in watch glasses filled with RNAlater Stabilization Solution. All instruments, surfaces, and tools were cleaned with 70% ethanol (Kopec) and RNase Away Decontamination Reagent (Thermo Scientific) prior to dissection. Forceps were also flamed prior to RNase Away treatment. Beetles were dissected and pooled into two tissue types: secretory lobe tissue and whole-body tissue without secretory lobes. Wings and elytra were stored separately as vouchers. Three biological replicates were collected for both taxa.

### RNA Extractions, Library Preparation, and Illumina Sequencing

*Platynus angustatus* and *Pterostichus moestus* samples were processed as described previously (Rork et al. 2021). Briefly, total RNA was extracted from both secretory lobe samples and whole-body samples using a standard TRIzol protocol and isolated using a DirectZol RNA MiniPrep Kit. Libraries were prepared using an Illumina Nextera Library Preparation kit. *P. angustatus* and *P. moestus* libraries were sequenced on an Illumina NovaSeq 6000 (two lanes, PE, 150 bp) at target reads depths of 25-30M/library.

### Computing Resources and Bioinformatics Software

All major bioinformatic analyses conducted as part of this study were carried out through the ROAR Collab supercomputer (RHEL 7/8) at The Pennsylvania State University unless otherwise noted. All bioinformatics software used was installed via the miniconda/anaconda package managers unless otherwise noted. A full list of software, packages, versions, and conda builds used in this work, as well as associated scripts and data files, can be found in SupplementaryData1.

### Quality Assessment of Reads and Trimming

The *Harpalus pensylvanicus* RNA-Seq data generated previously (see Rork et al. 2021) was reanalyzed as part of this study. Prior to quality assessment, FASTQ files generated from libraries split across multiple lanes (i.e. *Platynus angustatus* and *Pterostichus moestus*) were concatenated sample-wise. For all three species, initial FASTQC (v0.11.9) reports were generated for each sample and were manually inspected to flag any major data quality issues. After initial quality assessment, adapter content for every library was examined using BBTools’ (v38.90) bbduk.sh script using the adapters.fa file as a reference database (Bushnell 2014). Adapters were then trimmed using bbduk.sh and the literal adapter strings as references. The 3’ ends of reads were then trimmed to Q10 using the Phred algorithm as implemented by bbduk.sh. All species having been sequenced on two-color systems, poly-G tails were also trimmed. Reads containing ambiguous bases (Ns) were discarded, as were all reads shorter than 75 bp post-trimming. FASTQC reports were subsequently generated for each trimmed library to ensure detectable adapters, N-containing reads, and short reads were adequately removed without any trimming-induced degradation in library quality.

### Transcriptome Assembly and Quality Assessment

Transcriptomes were assembled *de novo* using Trinity RNA-Seq (v2.12) (Grabherr et al. 2011, Haas et al. 2013). Default settings were used for each assembly with a few noted exceptions. Read orientation (--SS_lib_type) was specified as “RF” for each species and minimum contig length (--min_contig_length) was set to 300 bp.

ExN50 values and transcript length statistics were generated using the “contig_ExN50_statistic.pl” and “TrinityStats.pl” scripts respectively. The “PtR” script was used to assess replicate-wise and sample-wise correlations within species. BUSCO was run on each transcriptome using default settings, the only exception being that analysis mode was set to “transcriptome” (Simão et al. 2015). The insect_odb9 database was used as the reference set of benchmark universal single-copy orthologs.

### Coding Sequence Prediction and Functional Annotation

Coding sequences were predicted *ab initio* from transcripts and were *in silico* translated to their respective polypeptides via TransDecoder (v5.5.0) (Hass et al. 2013). To determine homology of assembled transcripts and their predicted proteins BLASTX and BLASTP (v2.11.0) searches were run against the UniProtKB database using the full set of assembled transcripts and their predicted proteins as queries, respectively (Altschul et al. 1990). HMMScan (v3.3.2) searches were also run against the Pfam-A protein database using the full set of predicted proteins to assess protein domain content (Eddy 1998, Finn et al. 2011, Punta et al. 2012). For all BLAST and HMMScan searches, parameters were left as defaults except the e-value threshold set to 1e-10 (Altschul et al. 1990, Eddy 1998, Finn et al. 2011). Both the UniProtKB and Pfam databases were originally downloaded in March 2021 from the European Bioinformatics Institute’s FTP site. At the time of analysis, Trinotate was unable to be utilized for loading functional annotation results into databases (Hass et al. 2013). For this reason, a custom script was created to generate analogous databases, albeit without the typical eggNOG, KEGG, and GO information attached. A Trinotate (v3.2.2) database for each species was later generated when feasible.

### Read Quantification, Differential Gene Expression Analyses, and GO Enrichment

Transcript quantification was carried out using the Kallisto (v0.46.2) pseudoalignment-based strategy via the “align_and_estimate_abundance.pl” script included as part of Trinity RNA-Seq’s downstream analysis utilities (Bray et al. 2016). All settings were left as default. Transcripts and gene expression matrices were subsequently generated using the “abundance_estimates_to_matrix.pl” script, all settings again left as default.

Kallisto quantification results were used as inputs to the “run_DE_analysis.pl” script, used here to conduct differential gene expression analyses between tissues of each species (Bray et al. 2016). Specifically, contrasts were performed between secretory lobe (SL) samples and whole body (WB) samples in all species to identify genes significantly upregulated in the former relative to the latter. Voom was used to log2-transform gene-wise count data to log2-counts per million, and limma (v3.42.0) was used to identify differentially expressed genes from these data (Law et al. 2014, Ritchie et al. 2015). Genes were considered differentially upregulated in the secretory lobes at logFC > 4.0 and FDR < 0.05 (Benjamini & Hochberg 1995, Storey 2002). Note that this logFC cutoff is rather high compared to the usual cutoff of 1.5-2.0. This was done to only focus on the most upregulated genes in the secretory lobes, those which are putatively most important for their function. Notable genes logFC values < 4.0 are also discussed where appropriate.

GO enrichment analyses were run using GOSeq (v1.38.0) via the run_goseq.pl script, which was modified such that all GO Terms would be printed rather than only those with FDR < 0.05 (Benjamini & Hochberg 1995, The Gene Ontology Consortium 2000, Young et al. 2010). Lists of enriched and depleted GO terms for the secretory lobes relative to the whole bodies were generated for each species. The go_basics.obo file was downloaded manually, as the GO Consortium link changed between Trinity version releases (August 2021). Terms were classified as significantly enriched at FDR < 0.05. GO enrichment results were further summarized using the R package RRVGO (v1.4.4), which was not installed via miniconda3 (Sayols 2020).

### Phylogenetic Reconstruction of SSF and ABH Gene Families

Understanding the phylogenetic relationships between upregulated genes is necessary to assess the patterns underlying their evolution, their co-option, and their putative functions. Profile HMMs (pHMM) were downloaded for the sodium:solute symporter protein family (PF00474) and the alpha/beta hydrolase family (PF00561) from InterPro. The full set of coding and protein sequences from *Drosophila melanogaster* (GCF_000001215.4) and *Tribolium castaneum* (GCF_000002335.3) were downloaded from the RefSeq FTP server and the sequence headers shortened to species abbreviations and accessions. The coding and protein sequences of all three carabids, *D. melanogaster*, and *T. castaneum* were then concatenated into one multifasta file per sequence type. For the carabid protein sequences, all but the longest isoform per gene was removed.

Each pHMM was searched against the concatenated protein multifasta file using hmmsearch (HMMER v3.1b2) with an e-value reporting threshold of 1E-10. Accessions of hits were used to extract corresponding sequences from the concatenated coding and protein multifasta files (as coding and protein sequence accessions are identical in NCBI). Extracted coding sequences were aligned with MAFFT (i.e. not codon-aware) (v7.505) with the --auto parameter set, alignments were trimmed with trimAl (v1.4.rev15) with the --automated1 parameter set, and gap-rich sequences (comprised of >90% gap) were removed with the get_sequences_gaps_ratio.py script from trimAl (Capella-Gutiérrez et al. 2009, Katoh & Standley 2013).

Maximum-likelihood phylogenies were constructed for both gene families using IQ-Tree2 (v2.1.4-beta) with the following parameters set: -m MFP (standard model selection followed by tree inference) -B 5000 (5000 ultrafast bootstrap replicates) --merit BIC (BIC used to determine best-fit model of evolution) (Minh et al. 2020). Ten independent runs were carried out per gene family (--runs 10) and the topology with the highest likelihood was chosen. Resulting maximum-likelihood tree files were imported to R (v4.1.3) and the ggplot2 (v3.4.2) and ggtree (v3.2.1) packages were loaded to annotate the phylogenies (Paradis et al. 2004, Yu et al. 2017).

### Identification of Chemotype-specific Enriched GO Terms

To better assess which processes are important for chemical defense in a specific chemotype, we conducted multispecies combinatorial analyses to identify those GO terms commonly enriched in the secretory lobes of certain taxa. Specifically, we identified enriched GO terms exclusive to the secretory lobes of our formic acid producers and enriched GO terms exclusive to the secretory lobes of our methacrylic acid producer. This was done by piping lists of enriched GO terms of each species into successive of grep commands, at each step filtering out terms found in the non-target chemotype, thus leaving only terms shared by members of a chemotype at the end of the filtering process. To generate our list of formic acid-specific GO terms, for example, we first file grepped the list of secretory lobe-enriched GO IDs of *Harpalus pensylvanicus* against the list of secretory lobe-enriched GO IDs of *Platynus angustatus*. This gave us a subset of GO Terms enriched in both species’ secretory lobes. We then inverse grepped this list using the secretory lobe-enriched GO IDs of *P. moestus*, effectively removing from the initial list any secretory lobe-enriched GO IDs shared between “*H. pensylvanicus* AND *P. angustatus*” and *P. moestus*. This final list of GO IDs thus represented formic acid producer-specific secretory lobe-enriched GO IDs, which were then matched back to their respective GO Terms for interpretation. The same approach was used for *P. moestus*, only that inverse grep was exclusively used since we had no secondary methacrylic acid producer.

## RESULTS

### GC-MS Analysis of Pygidial Gland Defensive Chemicals

As described previously, we detected formic acid, undecane, and 2-tridecanone as the primary volatile components in the pygidial gland sprays of *Harpalus pensylvanicus*; an ensemble of compounds typical for this genus (Rork et al. 2021). The pygidial glands of *Platynus angustatus* also contained formic acid, undecane, and 2-tridecanone together with acetic acid, 2-tetradecanone, and 2-pentadecanone (Fig. 2a). To our knowledge, this is the fourth known instance of formic acid being produced by members of the genus *Platynus*, the second instance of 2-tridecanone detection, and the second of undecane detection (Kanehisa & Kawazu 1985). The remaining three compounds had not been detected in the pygidial gland defensive secretions of *Platynus* spp. to date (Schildknecht 1970, Kanehisa & Murase 1977, Kanehisa & Kawazu 1985). The pygidial gland secretions of *Pterostichus moestus* a total of five compounds were detected: acetic acid, propanoic acid, isobutyric acid, methacrylic acid, and tiglic acid (Fig. 2b-c). This is the twenty-fourth known instance of methacrylic and tiglic acids in *Pterostichus* spp., the second instance of acetic acid and 2-methylpropanoic acid detection, and the first of propanoic acid detection (Schildknecht 1970, Kanehisa & Murase 1977, Kanehisa & Kawazu 1982, Will et al. 2000).

**Figure 2a-c:**
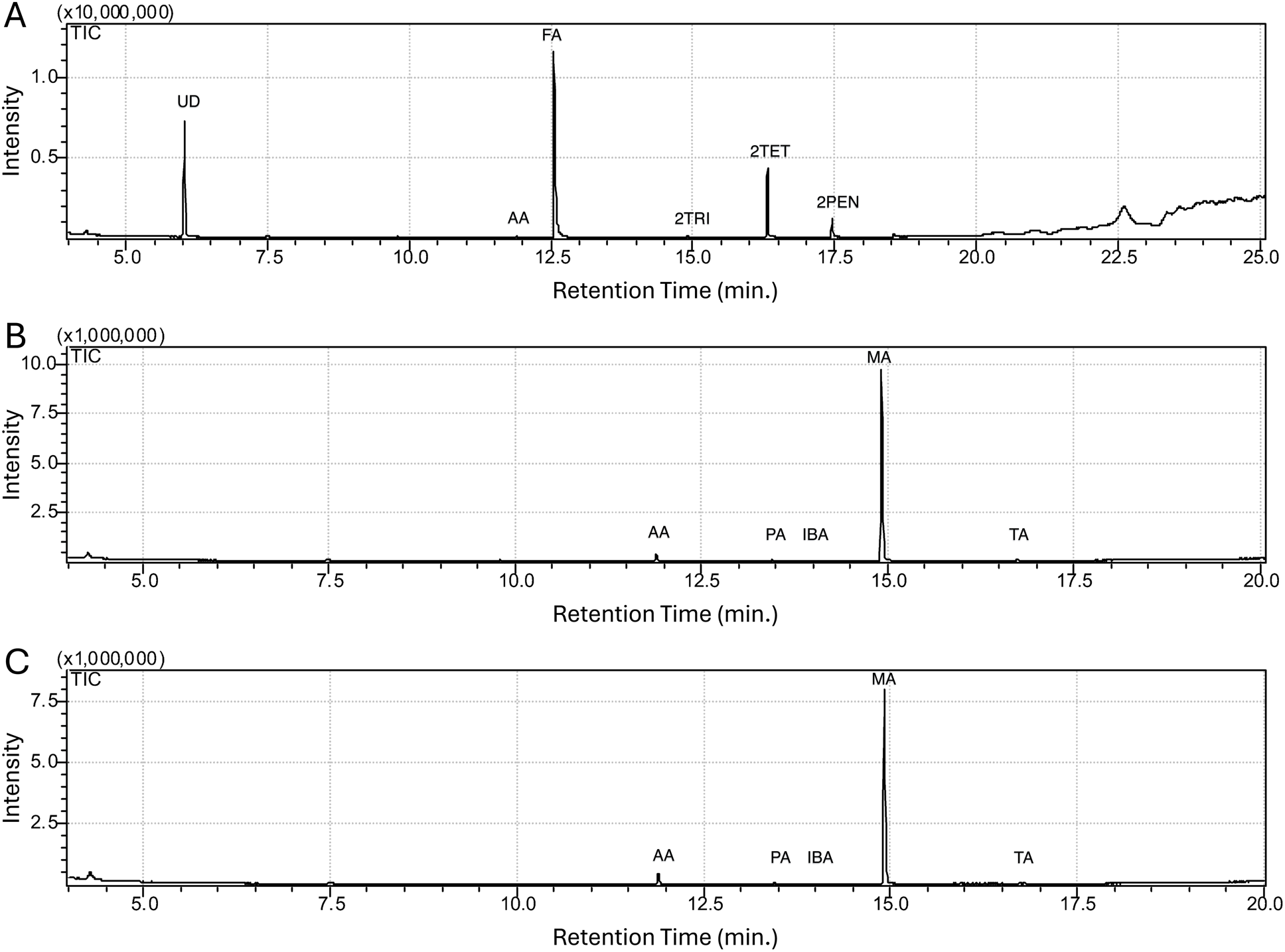
Gas chromatographs of the pygidial gland secretions of *Platynus angustatus* (a) and *Pterostichus moestus* (b: female, c: male). Peaks corresponding to organic acids are as follows: AA) Acetic acid, FA) Formic acid, PA) Propanoic acid, IBA) Isobutyric acid, MA) Methacrylic acid, TA) Tiglic acid. Peaks corresponding to hydrocarbons and ketones are as follows: UD) Undecane, 2TRI) 2-tridecanone, 2TET) 2-tetradecanone, 2PEN) 2-pentadecanone. Retention time (in minutes) is displayed along the x-axis and peak intensity is displayed along the y-axis.

### RNA-Seq Statistics

Due to the light trimming parameters, we discarded little data in terms of total reads across all samples (1.9%-2.6%). For *Harpalus pensylvanicus*, we removed approximately 5.7% of all bases, 13.2% of bases for *Platynus angustatus*, and 15.1% of bases for *Pterostichus moestus*. In these two species, the percent of reads containing adapters were 46.1% and 47.7% respectively. Nevertheless, due the much higher depth of sequencing for *P. angustatus* and *P. moestus* relative to *H. pensylvanicus*, these two species retained a greater total number of reads post-trimming than *H. pensylvanicus*. A full list of summary statistics for the raw and trimmed RNA-Seq data can be found in Supplementary Table 1.

### Transcriptome Assembly Statistics

All species’ transcriptomes were almost entirely complete according to BUSCO, ranging from 97.4% to 98.5%. In all cases, the majority of BUSCOs present were duplicated (64.1% to 92.8%). Based on the criterion by which BUSCO assesses a gene to be “duplicated” (i.e. more than one homologous sequence found in the target sequence set), this inflated duplicated value is due more to the assembly of multiple isoforms per gene rather than true duplication events within these beetles’ genomes (Simão et al. 2015).

Regarding assembly size, *Pterostichus moestus* had the smallest number of assembled transcripts (105,544) and the shortest total assembly length (123 Mb). *Harpalus pensylvanicus* had the largest assembly in terms of both number of assembled transcripts (163,238) and total length (167 Mb). GC content was similar for all species, ranging from 36.3% to 38.4%. E90N50 values ranged from 1,649 bp to 1,750 bp.

Between 44.8% and 47.7% of all transcripts were predicted to contain at least one coding sequence. Across all species, between 32.8% and 35.8% of transcripts received one or more blastx hits, between 63.9% and 66.1% of all predicted peptides received one or more blastp hits, and between 59.7% and 60.4% of all predicted peptides received one or more hmmscan hits.

Per species, the average pseudoalignment rate across all samples was between 68.0% and 81.9%, the lowest being in *Harpalus pensylvanicus* and the highest in *Pterostichus moestus*. Differential gene expression analyses revealed between 488 and 1,010 genes to be significantly upregulated in the secretory lobes of all species with *P. moestus* having the fewest upregulated genes and *H. pensylvanicus* having the most. A full list of summary statistics for the transcriptome assembly and all downstream analyses can be found in Supplementary Tables 2-5.

### Candidate Genes and Pathways Involved in Formate Biosynthesis

In the secretory lobes of both formic acid-producers, *Harpalus pensylvanicus* and *Platynus angustatus*, we found evidence for the upregulation of all three genes involved in the core of the folate cycle of C-1 metabolism (Fig. 3a, 4a). We define upregulation as a significant increase in gene expression in the secretory lobes relative to the rest of the body. Specifically, these are serine hydroxymethyltransferase (*SHMT*), bifunctional methylenetetrahydrofolate dehydrogenase/cyclohydrolase (*MTHFD2*), and trifunctional methylenetetrahydrofolate dehydrogenase/cyclohydrolase, formyltetrahydrofolate synthetase (*MTHFD1*) (Fox & Stover 2008, Brosnan & Brosnan 2016, Meiser et al 2018). In the secretory lobe-upregulated gene sets of both *H. pensylvanicus* and *P. angustatus*, we found a single upregulated copy of *SHMT*. In *H. pensylvanicus,* we found one upregulated copy of *MTHFD2* and two upregulated copies of *MTHFD1*, whereas *P. angustatus* has one upregulated copy each of *MTHFD2* and *MTHFD1*. Other non-upregulated copies of these genes were also found in *H. pensylvanicus* and *P. angustatus*. As discussed in previous work, it is unlikely that these multiple copies are truly independent genes, but rather are artifacts of *de novo* transcriptome assembly from heterozygous, pooled conspecifics (Rork et al. 2021). Some also appear to be bacterial, possibly those of secretory lobe endosymbionts (McManus et al. 2018). Not only are these three genes upregulated in the secretory lobes of both formic acid producers, but they are exclusively so. That is, they are not upregulated in the secretory lobes of *Pterostichus moestus* (Fig 4a).

**Figure 3a-b:**
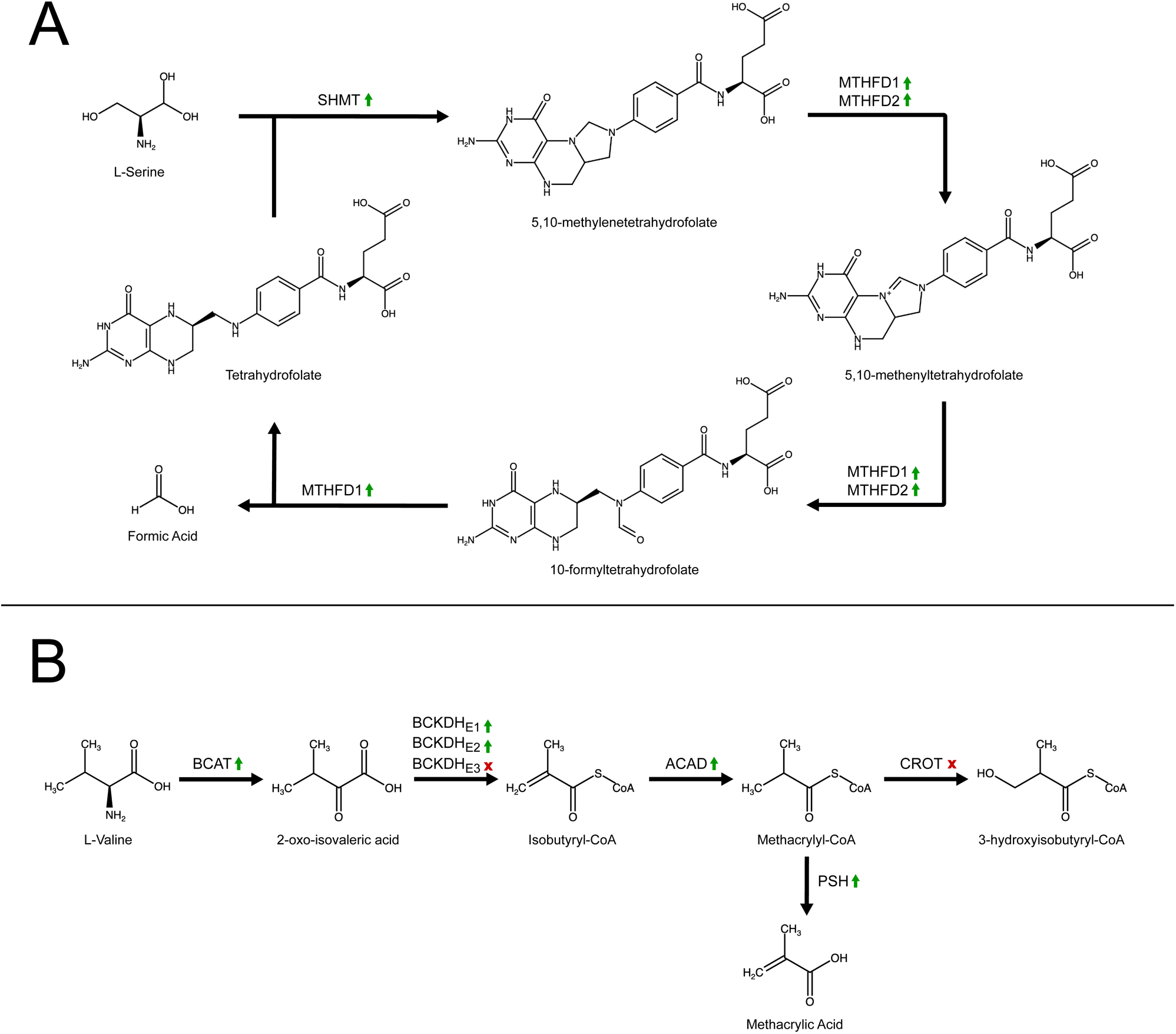
Simplified representations of the folate cycle of one-carbon metabolism (a) and the valine catabolic pathway (b). Compound names are shown below structures and gene/enzyme abbreviations above bolded black arrows. In (b), green arrows pointed upward to the right of gene/enzyme abbreviations indicate significant upregulation (logFC > 4, FDR < 0.05) in the secretory lobes of *H. pensylvanicus* and *P. angustatus*. In (b), the green arrows have the same meaning, but indicate upregulation in the secretory lobes of *P. moestus*. A red x to the right of the gene/enzyme abbreviations indicate that it is not significantly upregulated in *P. moestus* secretory lobes.

**Figure 4a-b:**
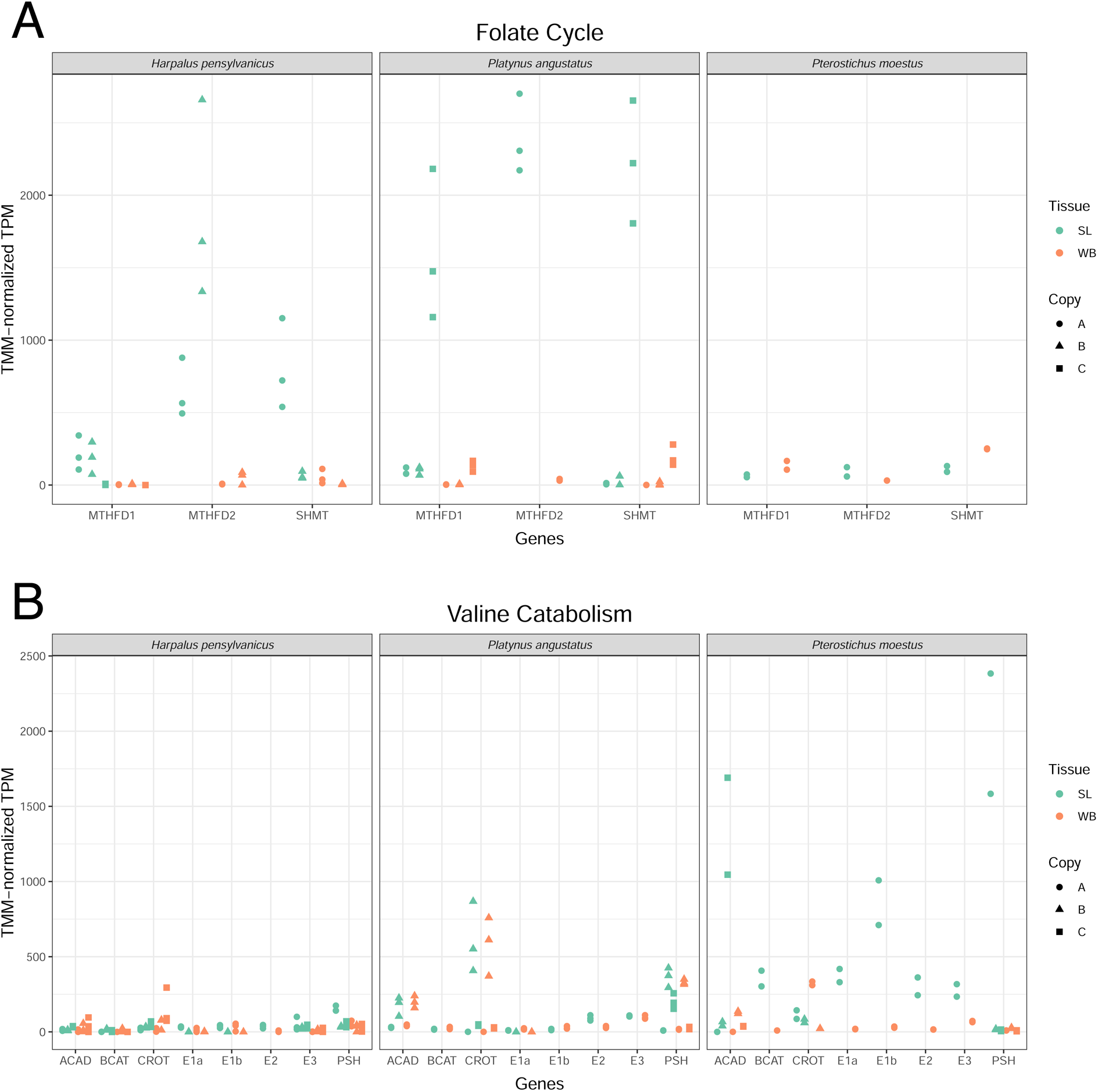
Expression of key genes involved in the folate cycle of one-carbon metabolism (a) and the valine catabolic pathway (b). Genes are listed along the x-axis whereas the y-axis measures TMM-normalized TPM gene expression values. Teal points (left side of the gene bins) represent secretory lobe samples and orange points (right side of the gene bins) represent whole body samples. Although all of these genes are likely single copy orthologs, for reasons discussed in this work, *de novo* assembly annotated multiple copies of most genes, represented here by differently shaped points. Up to the top three most highly expressed gene copies are displayed. This excluded no upregulated genes.

We also find evidence for the upregulation of one gene involved in the kynurenine pathway of tryptophan catabolism. Specifically, both *H. pensylvanicus* and *P. angustatus* upregulate kynurenine formamidase (*KFA*) in their secretory lobes, two copies being found in each upregulated gene set (Mehler & Knox 1950, Kanai et al. 2009, Han et al. 2012, Badawy 2017). We found no other genes involved in this pathway to be upregulated in either species’ secretory lobes. As with the three folate cycle genes, *KFA* upregulation in the secretory lobes is exclusive to the two formic acid-producers. Although we find evidence for the upregulation of 1,2-dihydroxy-3-keto-5-methylthiopentene dioxygenase of the methionine salvage pathway in *H. pensylvanicus* secretory lobes, we find no such evidence in *P. angustatus*, nor any evidence for the upregulation of any other gene in the methionine salvage pathway within the secretory lobes (Savarse et al. 1985, Sekowska et al. 2019). No other genes with established roles in formic acid biosynthesis, to our knowledge, were identified in either species’ upregulated gene sets.

There are several GO terms enriched exclusively in the secretory lobes of formic acid producers. Amongst those under the ontological category “Biological Process” are “Tetrahydrofolate metabolic process”, “Folic acid-containing compound metabolic process”, and “One-carbon metabolic process”, all which map to the core genes of the folate cycle. “Carboxylic acid metabolic process”, “Organic acid metabolic process”, and “Oxoacid metabolic process” are also exclusively enriched in these species’ secretory lobes, all of which again map to genes involved in the folate cycle, as well as the kynurenine pathway. Terms exclusively enriched under the “Molecular Function” category are even more specific, listing all primary functions of MTHFD1 and MTHFD2: “Methenyltetrahydrofolate cyclohydrolase activity”, “Methylenetetrahydrofolate dehydrogenase (NADP+) activity”, and “Formate-tetrahydrofolate ligase activity”. Although not specific to KFA, the exclusively enriched term “Hydrolase activity, acting on carbon-nitrogen (but not peptide) bonds” does refer to the activity of this enzyme according to the gene IDs mapped to it by GO-Seq.

### Candidate Genes and Pathways Involved in Methacrylate Biosynthesis

In the secretory lobes of *Pterostichus moestus*, we find all but one of the genes involved in the valine catabolic pathway, through the generation of methacrylyl-CoA, to be upregulated (Fig. 3b, 4b) (Lange et al. 2004, Wanders et al. 2010). Namely, these are branched-chain-amino-acid aminotransferase *(BCAT)*, the alpha and beta subunits of the E1 subunit of the branched-chain alpha-keto dehydrogenase complex (*E1a* and *E1b*), the E2 subunit of the branched-chain alpha-keto dehydrogenase complex (*E2*), and a short-chain specific acyl-CoA dehydrogenase (*ACAD*) (Brosnan & Brosnan 2006, Kochevenko et al. 2012). Only one copy of each gene is present in the upregulated gene set of this species’ secretory lobes. These genes are also exclusively upregulated in the secretory lobes of *P. moestus*, our sole methacrylic acid-producer; that is, they are not upregulated in the secretory lobes of *H. pensylvanicus* or *P. angustatus*. The one gene in this pathway not significantly upregulated in the secretory lobes of *P. moestus* is the E3 subunit of the branched-chain alpha-keto dehydrogenase complex (logFC = 2.44, FDR = 3.8E-3). Crotonase, which would normally convert methacrylyl-CoA to 3-Hydroxyisobutyryl-CoA, is similarly not significantly upregulated (logFC = 2.13, FDR = 5.5e-3).

To our knowledge, no known enzyme has been implicated in the conversion of methacrylyl-CoA to methacrylic acid. Presumably, such an enzyme would have thioesterase activity such that the CoA and acyl moieties could be cleaved and the latter hydroxylated to form the corresponding acid (Grevengoed et al. 2014). We do find one secretory lobe-upregulated gene which may have such a function: a probable serine hydrolase. Serine hydrolases comprise a large family of enzymes, many of which function as acyl-CoA hydrolases. Specifically, the serine hydrolase contains the same α/β hydrolase domain found in other acyl-CoA thioesterases, thus making it plausible that this enzyme could hydrolyze methacrylyl-CoA to methacrylic acid and CoA (Grevengoed et al. 2014). However, due to the lack of high-quality annotations for the genes in clade ABH-A of the larger α/β hydrolases gene family, we cannot make any generalizations on that basis alone (Fig. 5a-b). Clade ABH-B, which is relatively closely related to ABH-A and contains one upregulated sequence from *P. angustatus*, is comprised of lysophosphatidylserine lipases. These enzymes have acylglycerol lipase (esterase) and phospholipase (phosphoesterase) activity (Kelkar et al. 2019, Navia-Paldanius et al. 2012) (Fig. 5a, c). They have no known thioesterase activity, however, leaving the function of genes in clade ABH-A unknown.

**Figure 5a-b:**
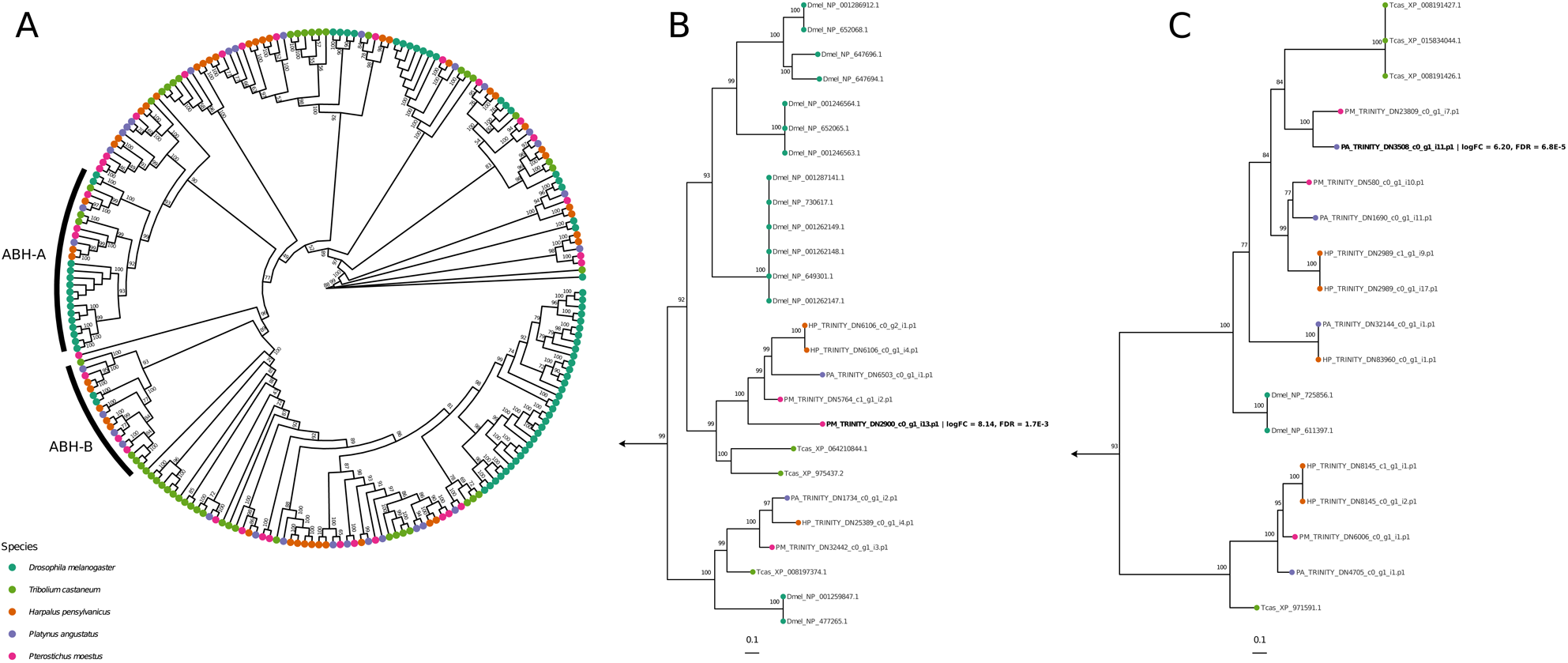
Maximum-likelihood phylogeny of the the α/β hydrolase (ABH) fold gene family, reconstructed via IQ-TREE2 from *Tribolium castaneum* (green), *Drosophila melanogaster* (teal), *Harpalus pensylvanicus* (orange), *Platynus angustatus* (purple), and *Pterostichus moestus* (magenta) sequences (a). Ultrafast bootstrap support values (5000 replicates) shown at nodes. Subclades ABH-A (b) and ABH-B (c) are annotated in the full phylogeny and shown separately.

There are also many related GO terms enriched exclusively in the secretory lobes of *P. moestus*. Under “Biological Process” are the closely related terms “branched-chain amino acid catabolic process” and “branched-chain amino acid metabolic process”, both of which describe the breakdown of branch-chain amino acids to their various acyl-CoA derivatives. Under “Cellular Component” are terms relating specifically to the BCKDH complex, namely “dihydrolipoyl dehydrogenase complex”, “mitochondrial alpha-ketoglutarate dehydrogenase complex”, “tricarboxylic acid cycle enzyme complex”, and “mitochondrial tricarboxylic acid cycle enzyme complex”. This suggests the enrichment of genes involved in such processes, all of which relate to L-valine catabolism and methacrylyl-CoA biosynthesis.

### Candidate genes involved in carboxylic acid transport

As important as the ability to synthesize defensive chemicals is to carabids, so too is their ability to secrete them from their secretory lobes into the lumens of the collecting ducts and reservoirs. In our three carboxylic acid producing taxa, this may be accomplished via monocarboxylate symporters, many of which belong to the sodium:solute symporter family (SSF) (Jung 2002).

Phylogenetic reconstruction of the sodium:solute symporter gene family suggests that members of two distinct clades may be important for monocarboxylate transport in the secretory lobes of *Harpalus pensylvanicus*, *Platynus angustatus*, and *Pterostichus moestus* (Fig. 6a-c). Here, we denote these clades as SSF-A and SSF-B. Both SSF-A and SSF-B contain gene family members from all three carabid species, but the genes upregulated in the secretory lobes of *H. pensylvanicus* and *P. angustatus* are found exclusively in SSF-A (98% UFBS) whereas the gene upregulated in the secretory lobes of *P. moestus* is exclusive to SSF-B (100% UFBS). Within SSF-A, the two upregulated *H. pensylvanicus* genes are sister (100% UFBS) and are themselves sister to two *P. angustatus* genes, one upregulated and one not (100% UFBS). This four-gene clade is nested within a clade containing a second upregulated *P. angustatus* gene (99% UFBS) (Fig. 6a-b). The one upregulated *P. moestus* sequence is upregulated in a distantly related clade (SSF-B) (Fig. 6a, c). The upregulation of likely orthologous (or at least closely homologous) sodium:solute symporters in the two formic acid producers and a relatively distantly-related sodium:solute symporter in the methacrylic acid producer suggests that if they are involved in the transport of the primary defensive chemicals into the secretory lobe lumens, they may transport different compounds. However, it is not clear from the theoretically better annotated yet functionally unvalidated *Drosophila* and *Tribolium* genes in these clades which classes, sizes, and polarities of solutes they are prone to transport.

**Figure 6a-c:**
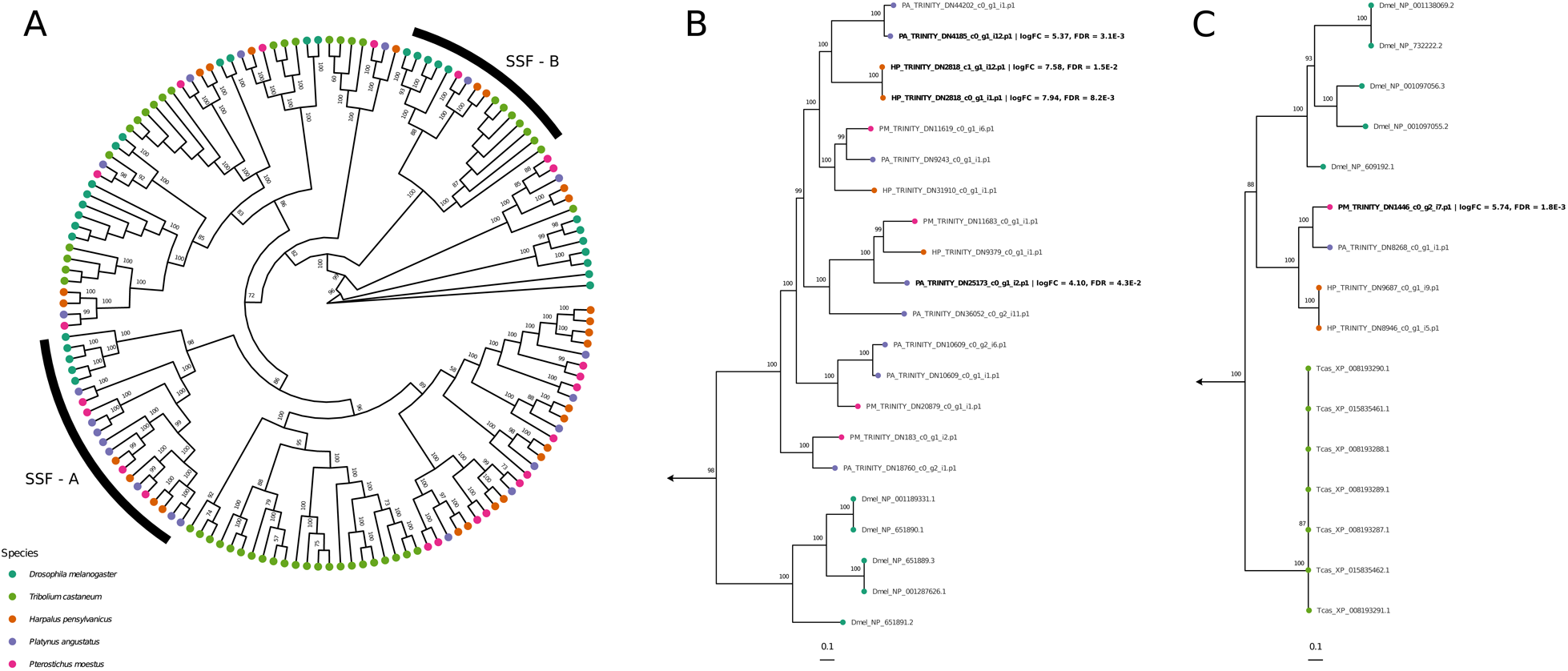
Maximum-likelihood phylogeny of the the sodium:solute symporter (SSF) gene family, reconstructed via IQ-TREE2 from *Tribolium castaneum* (green), *Drosophila melanogaster* (teal), *Harpalus pensylvanicus* (orange), *Platynus angustatus* (purple), and *Pterostichus moestus* (magenta) sequences (a). Ultrafast bootstrap support values (5000 replicates) shown at nodes. Subclades SSF-A (b) and SSF-B (c) are annotated in the full phylogeny and are shown separately.

## DISCUSSION

### Defensive-grade formic acid is likely biosynthesized similarly in *Harpalus* and *Platynus*

The results of our differential gene expression analyses and GO enrichment analyses suggest up to two pathways may be involved in defensive-grade formic acid biosynthesis in both *Harpalus pensylvanicus* and *Platynus angustatus*. The primary candidate pathway thought to be involved is the folate cycle of one-carbon metabolism, which has also been implicated in formic acid biosynthesis in ants (Hefetz & Blum 1978a, Hefetz & Blum 1978b, Rork et al. 2021). The core of this cycle is comprised of three enzymes: serine hydroxymethyltransferase, bifunctional methylenetetrahydrofolate dehydrogenase/cyclohydrolase, and trifunctional methylenetetrahydrofolate dehydrogenase/cyclohydrolase, formyltetrahydrofolate synthetase (Fox & Stover 2008, MacFarlane et al. 2009, Brosnan & Brosnan 2016). While tetrahydrofolate is a necessary input to the pathway, the one-carbon donor and precursor to formic acid itself is often L-serine (Hefetz & Blum 1978a, Hefetz & Blum 1978b). The stable intermediates in this pathway, 5,10-methylenetetrahydrofolate (5,10-mTHF) and 10-formyltetrahydrofolate (10-fTHF), are also intermediates in the biosynthesis of thymidylate and purines respectively (Hartman & Buchanan 1959, Carreras & Santi 1995, Ben-Sahra et al. 2016). When the former is reduced by methylenetetrahydrofolate reductase (MTHFR), the 5-methyltetrahydrofolate (5-mTHF) product is also utilized to regenerate methionine from homocysteine via methionine synthase (Zheng et al. 2019). Excess formate is typically exported from the cell into the extracellular space (as in carabid beetles secretory lobes and oxidative cancer cells) or is used in other metabolic processes (Meiser et al. 2016).

The second candidate pathway potentially involved in defensive-grade formic acid biosynthesis is the kynurenine pathway. This pathway begins by the oxidation of the imidazole ring of tryptophan, generating N-formylkynurenine in the process (Mehler & Knox 1950, Han et al. 2012, Badawy 2017). N-formylkynurenine is then hydrolyzed by kynurenine formamidase, cleaving off the formyl group as formate and generating L-kynurenine. L-kynurenine can then be used to synthesize a variety of compounds, most notably nicotinamide adenine dinucleotide (NAD) (Badawy 2017).

Of these two, we view it more likely that the folate cycle is the primary pathway responsible for defensive-grade formic acid biosynthesis for two primary reasons. The first is that L-serine is much less bioenergetically expensive than L-tryptophan, making it a more suitable precursor for the biosynthesis of relatively large quantities of formate (Moura et al. 2013, Barik 2020, Wu et al. 2020). Not only is L-tryptophan typically the least abundant amino acid in organisms, but animals cannot biosynthesize it (Moura et al. 2013). L-serine, on the other hand, is comparatively more abundant in organisms and can be biosynthesized by animals, typically from L-glycine or 3-phosphoglycerate (Moura et al. 2013, Wu et al. 2020). The second reason relates to the relative metabolic stress likely imposed by the upregulation of each pathway. By virtue of being a cycle, the only outputs of the folate cycle when it is primarily involved in biosynthesizing formate are formate, tetrahydrofolate, L-glycine, NAD(P)H, and ATP (Fox & Stover 2008). With formate being exported to the extracellular space, this leaves only an accumulation of L-glycine, tetrahydrofolate, NAD(P)H, and ATP. Tetrahydrofolate can be reused by the folate cycle and NAD(P)H and ATP are likely needed in abundance due to the high metabolic activity of the glands. On the other hand, the kynurenine pathway is acyclic, and thus would be producing a large quantity of comparatively specific metabolites that would need to serve some niche metabolic purpose or be degraded (Badawy 2017). Of course, it is possible that the products of the kynurenine pathway are important for secretory lobe function, but combined with the bioenergetic cost of using L-tryptophan as the primary precursor to formate biosynthesis, we view this as the less probable of the two options.

In our previous work, we also suggested the methionine salvage cycle as a potential albeit unlikely pathway for formate biosynthesis in *H. pensylvanicus*, due to its upregulation of 1,2-dihydroxy-3-keto-5-methylthiopentene dioxygenase in the secretory lobes (Rork et al. 2021). While this gene is still upregulated in our new *H. pensylvanicus* assembly, it is not in *P. angustatus*. While that alone is not reason enough to suggest it is unlikely to be involved in formate biosynthesis in *H. pensylvanicus*, the methionine salvage cycle shares some of the same conceptual issues as the kynurenine pathway and thus is not likely to be a major contributor in our view. It is also possible that it does have some limited involvement in formate biosynthesis in *H. pensylvanicus*, its role in *P. angustatus* perhaps just being even more limited if there is one.

It is important to note that while both *H. pensylvanicus* and *P. angustatus,* members of the same subfamily, seem to upregulate the same genes for formic acid biosynthesis in their secretory lobes, they likely did not converge upon this phenotype. Prior work supports the notion that the formic acid chemotype was derived from a common ancestor in these two genera (Ober & Heider 2010, Ober & Maddison 2008, Will et al. 2000). However, our data nevertheless can be seen as two independent data points supporting the hypothesis that the folate cycle and perhaps the kynurenine pathway are involved in defensive-grade formic acid biosynthesis in these taxa.

### Valine catabolism may underlie methacrylic acid biosynthesis in *Pterostichus*

The results of our differential gene expression and GO enrichment analyses for *Pterostichus moestus* suggest the involvement of the valine catabolic pathway in the biosynthesis of defensive grade methacrylic acid. This pathway begins with the transamination of L-valine to form 2-oxoisovaleric acid, which is then decarboxylated and covalently bound to coenzyme A to form isobutyryl-CoA (Brosnan & Brosnan 2006, Wanders et al. 2010). Isobutyryl-CoA is then reduced to methacrylyl-CoA, which can be further catabolized to ultimately form succinyl-CoA, an intermediate in the citric acid cycle (Wanders et al. 2010). Necessarily, an enzyme capable of hydrolyzing the thioester bond of methacrylyl-CoA would instead be capable of generating the final methacrylic acid product detected in the pygidial glands of *P. moestus*. We hypothesize that this enzyme is a serine hydrolase, some of which have been functionally characterized as having acyl-CoA ligase or thioesterase activity (Long & Cravatt 2011, Grevengoed et al. 2014). The probable serine hydrolase upregulated in the secretory lobes of *P. moestus* specifically belongs to the α/β-hydrolase (ABH) family which does contain members having thioesterase activity, possibly capable of hydrolyzing methacrylyl-CoA to methacrylic acid (Long & Cravatt 2011) (Fig. 5a-c). This of course remains speculative and would require functional characterization to confirm.

Interestingly, several other compounds present in the pygidial secretions of *P. moestus* are likely generated through the same metabolic module, the wider catabolism of branched-chain amino acids. L-valine, L-leucine, and L-isoleucine can all be reduced to isobutyryl-CoA, isovaleryl-CoA, and (S)-2-methylbutanoyl-CoA by the activity of BCAT and the BCKDH complex, respectively (Wanders et al. 2010, Brosnan & Brosnan 2006, Kochevenko et al. 2012). Through the activity of generalist short-chain acyl-CoA dehydrogenases or substrate-specific dehydrogenases, these molecules are further reduced to methacrylyl-CoA, 3-methylcrotonyl-CoA, and tiglyl-CoA respectively. Tiglyl-CoA is further converted to Propanoyl-CoA. Isobutyryl-CoA, methacrylyl-CoA, tiglyl-CoA, and propanoyl-CoA are all simply the acyl-CoA esters of the carboxylic acids isobutyric acid, methacrylic acid, tiglic acid, and propanoic acid, respectively (Fig. 2b-c). Throughout the branched-chain carboxylic acid-defended Carabidae, several other carboxylic acids such as crotonic acid, isovaleric acid, angelic acid, etc., are likely also produced via these three interconnected pathways (Moore & Wallbank 1968, Kanehisa & Murase 1977). The relationship between these pathways and the defensive chemistry of such Carabidae is further supported by the following evidence: the stable isotope D8-L-Valine is incorporated into methacrylic and isobutyric acids in *Scarites subterraneus*; [2,3,4,4-(2)H(4)]isoleucine is incorporated into tiglic, 2-methylbutyric, and ethacrylic acids in *Pterostichus californicus*; and L-[U-14C]-Valine, L-[U-14C]-Leucine, and L-[U-14C]-Isoleucine were incorporated into methacrylic acid, 3-methylcrotonic acid, and tiglic acid respectively in *Carabus yaconinus* (Adachi et al. 1985).

### Formic and methacrylic acid-producers may export defensive chemicals via sodium:solute symporters

Although it is ultimately uncertain which genes are involved in defensive chemical export from the secretory lobes in all taxa, we assess the most likely candidates in these carboxylic acid producers to belong to the sodium:solute symporter family. Specifically, this family includes proteins involved in monocarboxylate transport, including branched-chain fatty acids and formic acid (Jung 2002, Ganapathy et al. 2008, Moschen et al. 2012). We find it noteworthy that the two formic acid producers, *H. pensylvanicus* and *P. angustatus*, upregulate putatively orthologous or closely-related sodium:solute symporter homologs (Fig. 6a-b) while the methacrylic acid producer, *P. moestus*, upregulates a comparatively distantly related sodium:solute symporter homolog (Fig. 6a, c). Alternatives to monocarboxylate transporters that may be involved in the transport of formic and methacrylic acids across cell membranes include a variety of organic anion and cation transporters, many of which are members of the functionally diverse major facilitator superfamily (MFS) (Quistgaard et al. 2016). However, those few upregulated in the secretory lobes of *Platynus angustatus* generally seem unlikely to be involved in formate transport, but rather trehalose transport. This does not preclude the possibility of *Harpalus pensylvanicus* or *Pterostichus moestus* using MFS or other gene family members instead of or in conjunction with SSF gene family members or yet other transporters for defensive chemical export.

## CONCLUSIONS

The Carabidae are a taxonomically and biochemically diverse lineage whose chemical defense strategies have fascinated evolutionary biologists since the time of Darwin (Darwin 1846). Over the past several decades, considerable efforts have been undertaken to characterize the diversity of these defensive chemicals, leading to the accrual of hundreds of known compounds from carboxylic acids to phenolics, from sulfides to cyanide and more. While there has been much interest in how ground beetles (and insects more broadly) synthesize their defensive compounds, few pathways have been deduced and only for a handful of taxa. Here, we provide transcriptomic evidence suggesting likely pathways for the biosynthesis of the two most common defensive chemicals across the Carabidae, formic acid and methacrylic acid.

This work builds upon prior research suggesting that defensive-grade formic acid is biosynthesized by insects from serine via the folate cycle and that defensive-grade methacrylic acid is biosynthesized in Carabidae through the catabolism of valine. We also provide evidence for a pathway not previously suggested to play a major role in formic acid biosynthesis in insects, the kynurenine pathway.

## ACKNOWLEDGEMENTS

We would like to thank Yuka Imamura and Teodora Orendovici of the Penn State College of Medicine Genome Sciences Facility for their assistance preparing and sequencing our cDNA libraries at The Pennsylvania State University. We thank Wendy Moore for her helpful discussion on technical aspects of this research as well as the composition of this manuscript. This material is based upon work supported by the National Science Foundation under Grant No. DEB 1556898 to Athula Attygalle, DEB 1762760 to Tanya Renner, and DBI 2109735 to Adam Rork. Any opinions, findings, conclusions, and recommendations expressed in this material are those of the authors and do not necessarily reflect the views of the National Science Foundation. This work is supported by the USDA National Institute of Food and Agriculture and Hatch Appropriations under Project #PEN04974 and Accession #7006543. This work was also supported by The Coleopterists Society’s Graduate Student Research Enhancement Award to Adam Rork. The findings and conclusions of this work do not necessarily reflect the views of The Coleopterists Society.

## ETHICS STATEMENT

As all beetles were collected on property of The Pennsylvania State University, permits were not needed.

## DATA ACCESSIBILITY STATEMENT

All RNA-Seq data generated in this study and Rork et al. 2021 are available through the NCBI Sequence Read Archive (Bioproject: PRJNA705215). All quality assessment statistics, gene expression data, functional enrichment data, phylogenetics data, scripts, etc., are available in the SupplementaryData directory hosted on Penn State ScholarSphere (https://scholarsphere.psu.edu/resources/bf571e7d-95f7-40b3-b507-69ed5b971e97).

## AUTHOR CONTRIBUTIONS STATEMENT

A.M.R., S.X., A.A., and T.R. conceptualized the design of the project, interpreted results, created figures, and wrote the manuscript. A.M.R. collected specimens, conducted molecular biology, and bioinformatic analyses. S.X. conducted chemical analyses. A.M.R., A.A., and T.R. obtained funding. All authors read and approved the final manuscript.

